# Ultrasound-mediated blood-brain barrier disruption improves anti-pyroglutamate3 Aβ antibody efficacy and enhances phagocyte infiltration into brain in aged Alzheimer’s disease-like mice

**DOI:** 10.1101/2021.01.15.426806

**Authors:** Qiaoqiao Shi, Tao Sun, Yongzhi Zhang, Chanikarn Power, Camilla Hoesch, Shawna Antonelli, Maren K. Schroeder, Barbara J. Caldarone, Nadine Taudte, Mathias Schenk, Thore Hettmann, Stephan Schilling, Nathan J. McDannold, Cynthia A. Lemere

**Author notes:** Corresponding authors: Cynthia A. Lemere, Ph.D., and Nathan J. McDannold, Ph.D.,. Co-first authors (Qiaoqiao Shi, Tao Sun). Co-senior authors (Nathan J. McDannold, Cynthia A. Lemere).

## Abstract

Pyroglutamate-3 amyloid-β (pGlu3 Aβ) is an N-terminally modified, toxic form of amyloid-β that is present in cerebral amyloid plaques and vascular deposits. Using the Fc-competent murine anti-pGlu3 Aβ monoclonal antibody (mAb), 07/2a, we present here a nonpharmacological approach using focused ultrasound (FUS) with intravenous (i.v.) injection of microbubbles (MB) to facilitate i.v. delivery of the 07/2a mAb across the blood brain barrier (BBB) in order to improve Aβ removal and restore memory in aged APP/PS1 mice, an Alzheimer’s disease (AD)-like model of amyloidogenesis.

Compared to sham-treated controls, aged APP/PS1 mice treated with 07/2a immediately prior to FUS-mediated BBB disruption (mAb + FUS-BBBD combination treatment) showed significantly better spatial learning and memory in the Water T Maze. FUS-BBBD treatment alone improved contextual fear learning and memory in aged WT and APP/PS1 mice, respectively. APP/PS1 mice given the combination treatment had reduced Aβ42 and pGlu3 Aβ hippocampal plaque burden compared to PBS-treated APP/PS1 mice.

Hippocampal synaptic puncta density and synaptosomal synaptic protein levels were also higher in APP/PS1 mice treated with 07/2a just prior to BBB disruption. Increased Iba-1^+^ microglia were observed in the hippocampi of AD mice treated with 07/2a with and without FUS-BBBD, and APP/PS1 mice that received hippocampal BBB disruption and 07/2a showed increased Ly6G^+^ monocytes in hippocampal CA3. FUS-induced BBB disruption did not increase the incidence of microhemorrhage in mice with or without 07/2a mAb treatment.

Our findings suggest that FUS is useful tool that may enhance delivery of an anti-pGlu3 Aβ mAb for immunotherapy. FUS-mediated BBB disruption in combination with the 07/2a mAb also appears to facilitate monocyte infiltration in this AD model. Overall, these effects resulted in greater sparing of synapses and improved cognitive function without causing overt damage, suggesting the possibility of FUS as a noninvasive method to increase the therapeutic efficacy in AD patients.

## Introduction

Alzheimer’s disease (AD), the most common form of dementia worldwide, is defined pathologically by the presence extracellular amyloid-β (Aβ) plaques and intracellular neurofibrillary tangles containing hyperphosphorylated tau in brain [1]. While Aβ peptide occurs as a monomer under normal conditions, small neurotoxic aggregates, known as soluble Aβ oligomers, accumulate and aggregate further to form amyloid plaques [1, 2]. Increased Aβ production and reduced clearance are thought to underly the gradual accumulation of aggregated Aβ in AD brain [3]. Passive immunization approaches and anti-Aβ immunotherapy have been used for boosting Aβ clearance in mouse models and humans. However, limited delivery of antibodies, systematic side effects due to the off-target effects, and high cost have been handicaps for effective usage of these strategies in clinical trials [4, 5]. One of the main obstacles of effective drug delivery to the central nervous system is the blood brain barrier (BBB), a semipermeable barrier comprised of endothelial cells, astrocytes and pericytes, that protects the brain by blocking entry of circulating toxins [6]. Unfortunately, the BBB also limits CNS penetration of drugs into brain [7, 8].

Here, we investigated an alternative approach to conventional anti-Aβ immunotherapy that facilitates enhanced and spatially targeted modulation of Aβ clearance, in order to establish a more precise and controlled treatment paradigm for AD. To enhance the antibody delivery and Aβ clearance in a targeted manner, we used microbubble (MB)-enhanced focused ultrasound (FUS) to open the blood-brain barrier (BBB) noninvasively, reversibly and repeatedly [9, 10]. In this process, intravenously injected preformed microbubbles concentrate mechanical forces and stresses onto the vasculature. These mechanical effects result in an increased permeability of the BBB that lasts for several hours. MB-enhanced FUS has been reported previously to enable delivery of exogenous anti-Aβ antibodies [11, 12] and a GSK-3 inhibitor [13] in AD mouse models and reduced plaque burden. Other studies have shown that ultrasound-induced BBB opening itself improved Aβ clearance and behavior in 7-month-old [14], 1-year-old [15], and 2-year-old [16] AD-like mouse models. Increased neuronal plasticity with more newborn neurons [14] and microglial activation [15, 17] were also observed. A clinical trial on MB-enhanced FUS (alone) in AD patients has begun [18].

Amyloid-Related Imaging Abnormalities (ARIA) are transient vascular adverse events, including vasogenic edema and microhemorrhage, that have been observed in anti-amyloid immunotherapy clinical trials, especially in individuals with pre-existing amyloid deposition in blood vessel walls known as congophilic amyloid angiopathy (CAA). It remains unclear whether FUS-mediated enhanced delivery of an anti-Aβ monoclonal antibody in aged AD transgenic mice with both plaque and vascular amyloid will be effective or if increasing antibody delivery to the brain will augment the incidence of microhemorrhage. In addition, the potential role of FUS in sparing synapse loss, a main and early pathological signature of cognitive decline in AD, is not well understood. We hypothesized that combining MB-mediated FUS with anti-Aβ immunotherapy may further enhance the immunotherapeutic effects via increased and localized passive delivery of immunotherapy agents as well as the recruitment and activation of immune cells which may facilitate plaque removal.

Pyroglutamate-3 Aβ (pGlu3 Aβ) is a pathological, toxic form of amyloid-β that is present in many plaques and vascular amyloid in human brain and AD-like amyloid transgenic (Tg) mouse models [19]. We obtained a murine anti-pGlu3 amyloid-β IgG2a mAb (07/2a) from Vivoryon Therapeutics AG (Halle, Germany) for this study. We previously characterized and determined *in vivo* and *ex vivo* the pathological efficacy and *in vivo* behavioral improvement using the IgG2a mAb, 07/2a, and its IgG1 version, 07/1, in APP/PS1dE9 Tg mice, a model of Alzheimer’s disease amyloidosis [20-23]. Here, we investigated whether FUS-mediated BBB disruption combined with an anti-pGlu3 Aβ mAb, 07/2a, can improve both clearance of amyloid plaques and cognition in aged APP/PS1dE9 mice without increasing the incidence of microhemorrhage. Additionally, pathological assessments including synapse density, monocyte infiltration, and the presence of plaque-associated gliosis were performed to elucidate potential mechanisms of the combination therapy effects.

## Results

### FUS system and the reliable disruption of BBB

A system to perform FUS-induced BBB disruption in mice without MRI guidance was assembled in a non-barrier facility where the animals undergo behavior testing (Fig. 1a). An 835 kHz FUS transducer was used to apply pulsed sonications (10 ms bursts at 2 Hz for 100 s) on two hippocampal targets in each hemisphere in conjunction with 100 µl/kg i.v. injections of Optison MBs. We conducted a pilot study to confirm that we could reliably target the hippocampus with this system. This confirmation was made with fluorescent images of Trypan blue delivery (Fig. 1c). After satisfactory results were achieved, APP/PS1dE9 mice were treated with i.v. infusion of pGlu3 Aβ mAb alone or before FUS-induced BBB disruption (FUS-BBBD) in three weekly sessions, followed by behavior tests, euthanasia and tissue harvest (Fig. 1b). Control animals (WT and Tg) received either FUS-BBBD alone or treatment with PBS by i.v. infusion.

**Fig 1.**
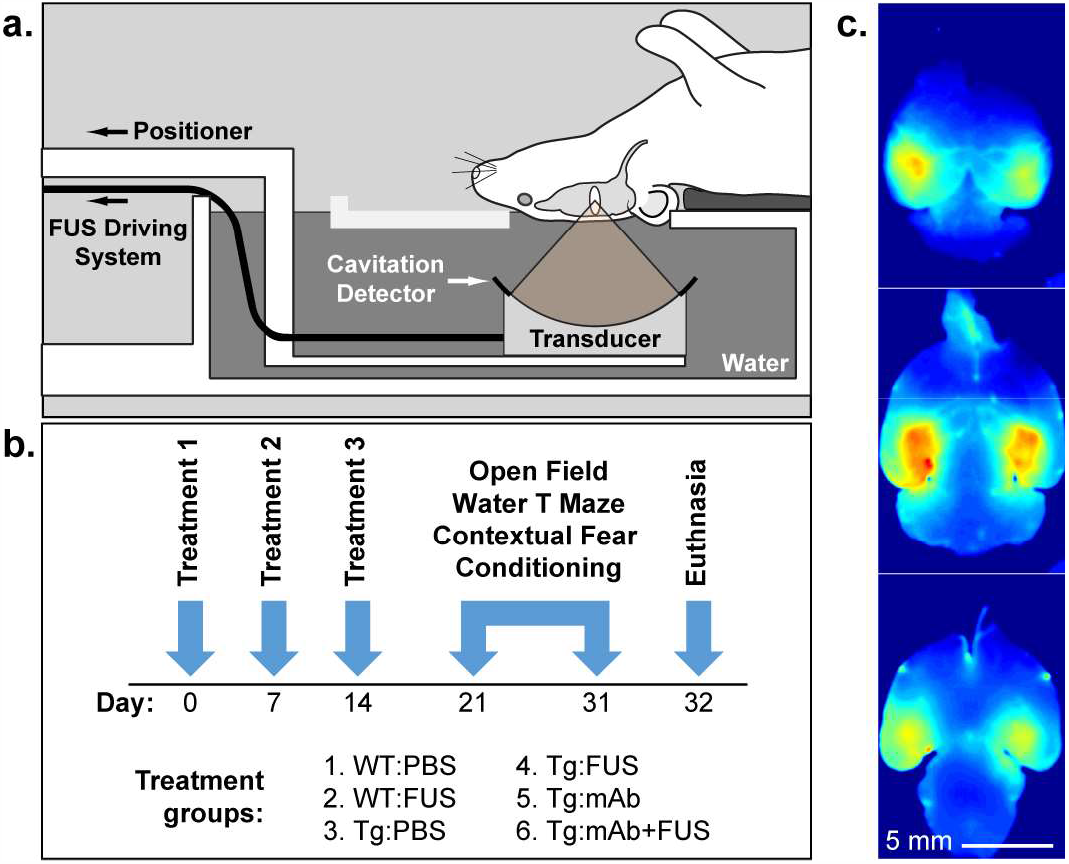
Treatment paradigm and fluorescent imaging of bilateral Trypan Blue delivery to the hippocampus. **a**. Male APP/PS1 mice at 16 months of age or wildtype (WT) age-matched male C57BL/6J mice were placed supine above the FUS transducer, which was positioned so that the focus was centered on the hippocampus. **b**. The mice were divided into 6 groups that received treatment with intravenous PBS, intravenous anti-pGlu3 Aβ mAb (07/2a), or intravenous anti-pGlu3 Aβ mAb immediately before BBB disruption via intravenous MB infusion and FUS-mediated BBBD. Mice were treated weekly for 3 weeks. Behavioral testing started 1 week after the final treatment and lasted 11 days, after which mice were sacrificed and tissue was collected for analysis. **c**. Fluorescence imaging of Trypan Blue delivery in a mouse brain after BBB disruption via MB-enhanced FUS. Two targets were sonicated in the hippocampus in each hemisphere in a pilot study to confirm the targeting accuracy of the system. After euthanasia and transcardial perfusion, the brain was cut into 2 mm slabs. The BBB disruption covered the hippocampus bilaterally and extended the full dorsal/ventral axis of the brain.

### Slowing of cognitive decline in APP/PS1 mice with combined anti-pGlu3 Aβ mAb and FUS-BBBD treatment

Hyperactivity in the Open Field test and impaired learning and memory have been reported in aged APP/PS1 mice compared with non-transgenic mice [21, 22, 24-26]. To determine whether treatment with an anti-pGlu3 Aβ mAb alone or combined with FUS-BBBD affects age-related cognitive decline, we assessed learning and memory in 16-month-old WT mice treated with PBS (WT:PBS), WT mice with FUS-BBBD (WT:FUS), APP/PS1 mice with PBS (Tg:PBS) or FUS alone (Tg:FUS), APP/PS1 mice treated with 07/2a mAb (Tg:mAb), and APP/PS1 mice treated an 07/2a combined with FUS-BBBD (Tg:mAb+FUS). We used the Water T Maze (WTM) test, which measures spatial learning and memory during the animal’s discovery and memory of the platform location [27]. Mice in the Tg:PBS group showed significant impairment in finding the platform over days 2-8 of testing compared to the mice in the WT:PBS group (*p* < 0.05; Fig. 2a). The percentage of these mice that reached a predetermined criterion (≥80% correct choices two days in a row) by Day 8, indicated that the Tg:PBS mice were unable to learn the task (Fig. 2b). Hence, the reversal test was not conducted in this study. The mice in the Tg:mAb+FUS group learned the location of the hidden platform more quickly than the Tg:PBS mice on days 5-8, and their performance was better than that of the mice in the Tg:mAb and Tg:FUS groups on day 5 (*p* < 0.05; Fig. 2a). Acquisition performance (i.e. learning and remembering the platform location) in Tg:mAb+FUS mice was similar to that of mice in the WT:PBS and WT:FUS groups, and it was better than in the mice in the Tg:mAb group (day 5, *p* < 0.05, Fig. 2a). By day 8, mice in the Tg:mAb+FUS group performed significantly better than Tg:PBS mice (*p* < 0.05) while mice in the Tg:mAb group showed a strong trend for improvement (*p* = 0.056) after 3 weekly treatments. Overall, these results suggest that enhanced delivery of the 07/2a anti-pGlu3 Aβ mAb to the hippocampus via FUS-BBBD enhanced spatial learning and memory in these aged APP/PS1 mice.

**Fig 2.**
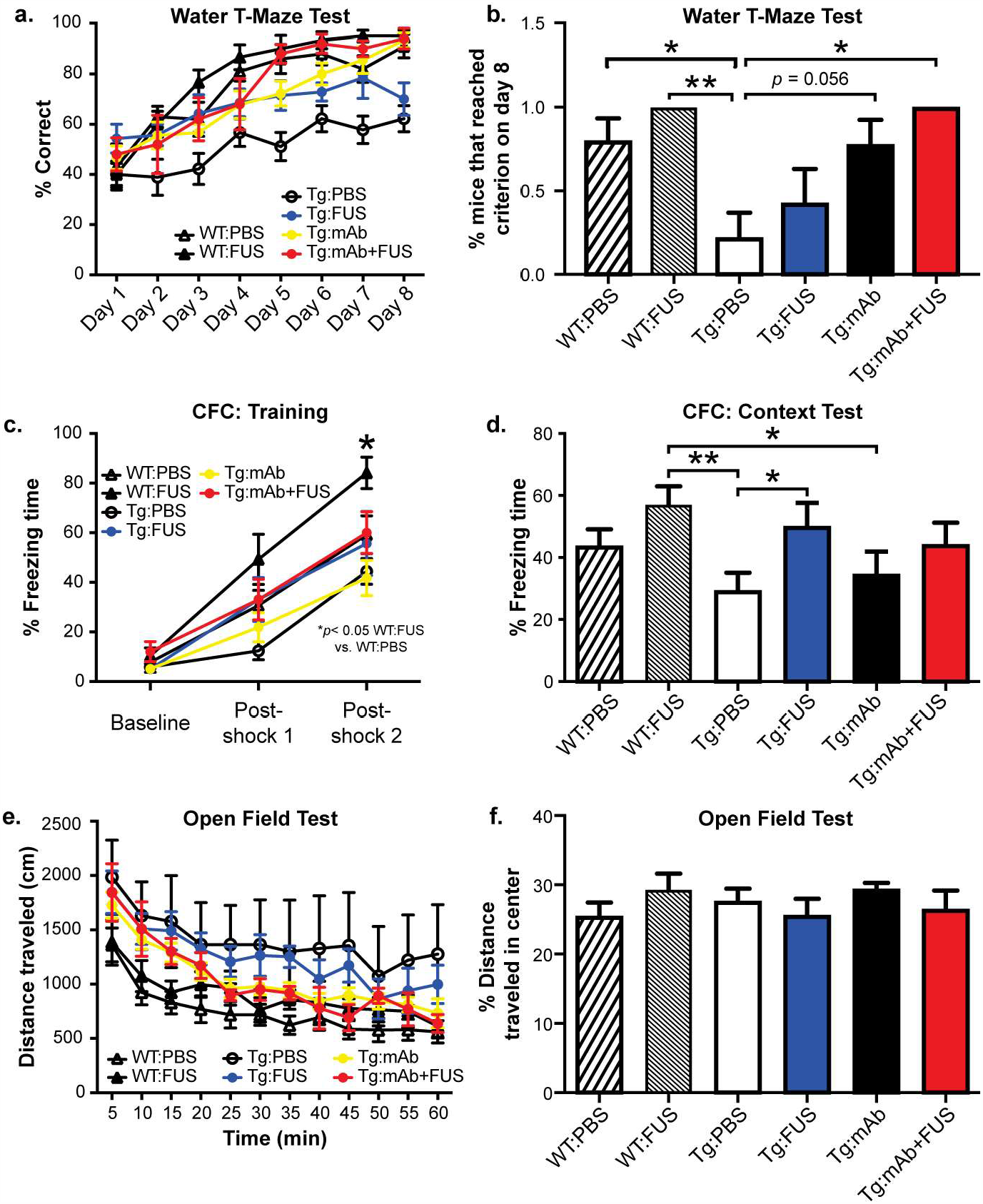
Behavior tests in WT and Tg mice with or without FUS and pGlu3 Aβ mAb, 07/2a. **a**. Tg:PBS mice were significantly impaired in the Water T Maze (WTM) test compared to WT:PBS mice (*p* < 0.001) at 16 months of age. The Tg:FUS, Tg:mAb, Tg:mAb+FUS groups showed significantly-improved learning and memory (days 5-8 of WTM acquisition) in the WTM compared to the Tg:PBS group (*p <* 0.05 for Tg:FUS vs Tg:PBS on Day 6; *p < 0*.*01 for Tg:mAb+FUS vs Tg:PBS on Day 5-8; p <0*.*01 for Tg:mAb vs Tg:PBS on Day 5-8)*. The scores in these three treatment groups were similar to the performance of WT mice treated with PBS or FUS. **b**. Percent of mice that reached a pre-defined criterion (≥ 80% correct choices on 2 consecutive days) in the WTM test on Day 8 of WTM testing. A higher percentage of mice reached this criterion in the Tg:mAb+FUS group compared to Tg:PBS (* *p* < 0.05), indicating improved performance. **c-d**. In the Context Fear Conditioning test (CFC), WT mice treated with FUS alone spent significantly more time freezing than PBS-treated WT mice during training on day 1 (c), indicating improved fear learning (\**p <* 0.05). Tg mice treated with FUS alone spent more time freezing on day 2 compared to Tg:PBS (d), suggesting improved fear memory (\**p <* 0.05). **e-f**. In the Open Field test, Tg:PBS mice showed increased locomotor activity compared to the WT:PBS animals (e), suggesting hyperactivity (\**p <* 0.05). No other significant differences were seen among the groups in distanced traveled (e) or between any groups in the percent distance traveled in the center (a criterion for anxiety behavior) (f) (WT:PBS, n=13; WT:FUS, n=8; Tg:PBS, n=9; Tg:FUS, n=7; Tg:mAb, n=9; Tg:mAb+FUS, n=6). Tests were assessed using one-way ANOVA followed by Fisher’s Protected Least Signficant Difference post-hoc test.

In the Contextual Fear Conditioning test (CFC), the post-shock freezing time in the mice in the WT:FUS group was significantly longer during the training period (fear learning) compared to mice in the WT:PBS group (*p* < 0.05; Fig. 2c). When placed back into the box the next day, the mice in the Tg:FUS group had a longer freezing time in the absence of a shock (fear memory) compared the mice in the Tg:PBS group (*p* < 0.05; Fig. 2d). In addition, mice in the WT:FUS group froze longer than mice in the Tg:PBS (*p* < 0.01) and Tg:mAb (*p* < 0.05) groups. Increased freezing time in the CFC training and context test suggests that FUS treatment alone improved fear learning in WT mice and fear memory in Tg mice, or alternatively, may have increased the sensitivity or overall fear response to the small electric shock.

Similar to our previous reports [21, 22] we found that mice in the Tg:PBS group traveled a significantly greater distance in the Open Field test compared to mice in the WT:PBS group (Fig. 2e), suggesting increased locomotor activity. Overall, no significant differences were observed between the mice in the Tg:FUS, Tg:mAb and Tg:mAb+FUS groups and those in the WT:PBS or WT:FUS groups, indicating that FUS-BBBD, anti-pGlu3 Aβ mAb treatment and their combination attenuated hyperactivity in the APP/PS1 mice (Fig. 2e). In addition, none of the treatment groups showed any differences in distance traveled in the center in the Open Field test (Fig. 2f), suggesting the treatments did not affect anxiolytic-like behavior.

### Decreased hippocampal plaque deposition in APP/PS1 mice co-treated with anti-pGlu3 Aβ mAb and FUS-BBBD

To determine whether the memory improvement we observed in the APP/PS1 mice with the combination of FUS-BBBD and 07/2 treatment was associated with decreased Aβ in the sonicated hippocampal targets, we examined the Aβ plaque load in the hippocampus and in the non-sonicated pre-frontal cortex (PFC) (Fig. 3). A significant decrease in Aβx-42 plaque deposition in the hippocampus was found in the Tg:mAb+FUS mice (*p* < 0.05, Fig. 3a,b). No significant differences in Aβx-42 plaque burden were observed between mice treated with PBS, 07/2a alone, or FUS alone (Fig. 3a-e). In addition, no significant difference was observed between any group in Aβx-42 plaque deposition in the PFC or in Aβx-40 plaque deposition in the hippocampus or PFC. PyroGlu3 Aβ plaque deposition in the hippocampus was reduced in both the Tg:mAb and the Tg:mAb+FUS groups but only reached significance in the Tg:mAb+FUS group compared to the PBS-treated mice (Fig. 3f, g). No significant differences in pGlu3 Aβ plaque deposition were seen in PFC (Fig. 3h). The lack of a reduction in plaque burden in the PFC indicates that the effect was localized to the sonicated area. Immunostaining with an anti-mouse IgG2a secondary antibody alone to detect 07/2a in mouse brain sections revealed similar patterns between groups (data not shown).

**Fig 3.**
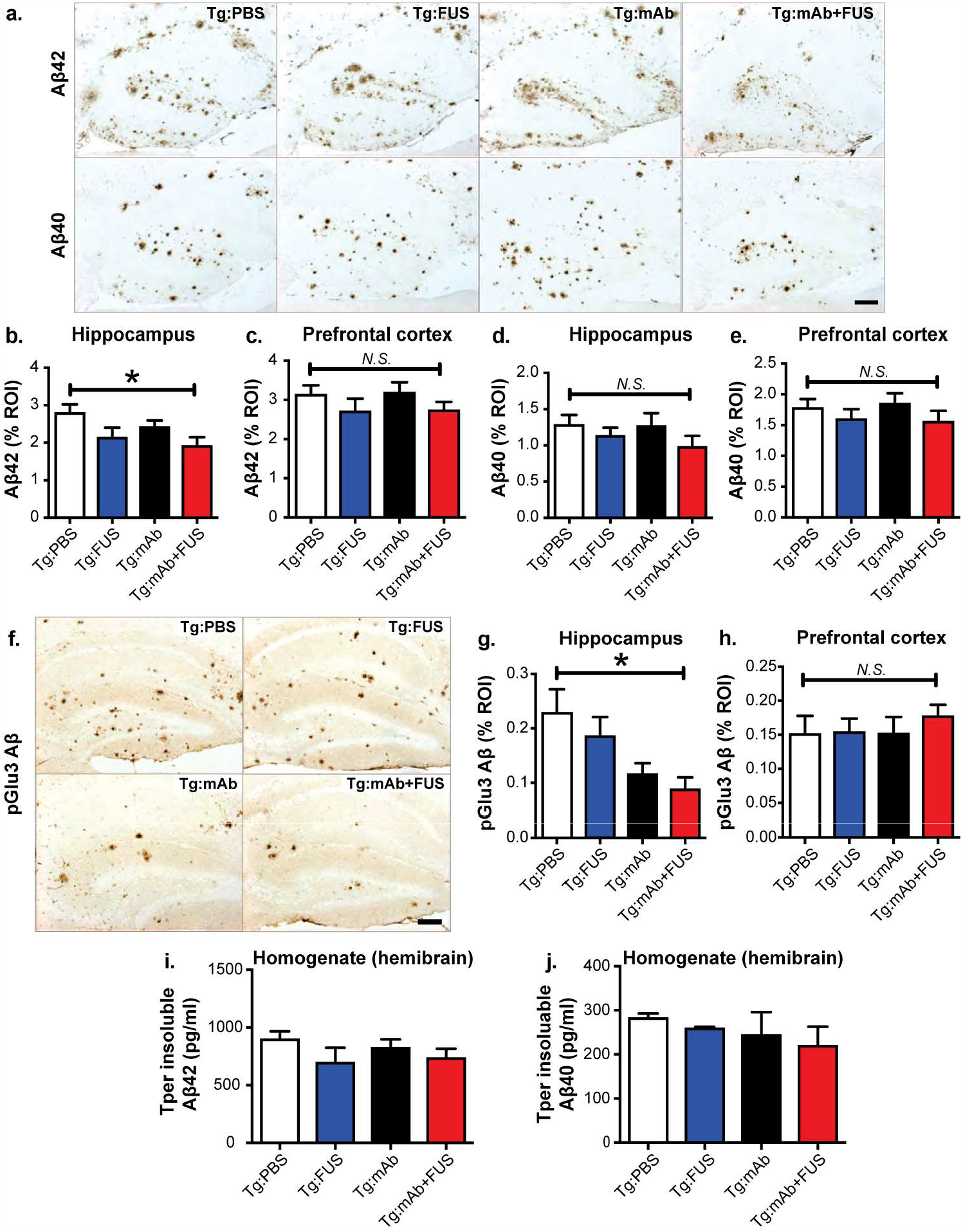
Aβ plaque load in 16 month-old Tg mice treated with PBS, FUS, anti-pGlu3 Aβ mAb, or a combination of mAb and FUS. **a-e**. Aβ42 immunoreactivity within the region-of-interest (ROI) showed reduced plaque burden in mice in the Tg:mAb+FUS group vs. Tg:PBS group (a, top row). No such reduction was observed in sections stained for Aβ40 immunoreactivity (a, bottom row). Quantification of the area of Aβ42-positive plaques showed a significant (*\*p < 0*.*05)* lowering in the hippocampus (b), but not the prefrontal cortex (PFC, c), in mice in the Tg:mAb+FUS group compared to mice in the Tg:PBS group. No significant difference among treatment groups was observed in the area of Aβ40 plaques in either hippocampus (d) or PFC (e). Scale bar = 100 μm. **f-h**. pGlu3 Aβ immunoreactivity in sections stained with K17 showed significantly decreased plaque load in the hippocampus RO1 in mice in the Tg:mAb+FUS group compared mice in the Tg:PBS group (f, g) (\**p <* 0.05). No significant difference was found in pGlu3-Aβ immunoreactivities in the PFC (h). Scale bar = 100 μm. **i-j**. No significant differences were seen in Aβ42 and Aβ40 levels by ELISA in whole hemibrain homogenates. (One-way ANOVA followed by Bonferroni test, WT:PBS, n=13; WT:FUS, n=8; Tg:PBS, n=9; Tg:FUS, n=7; Tg:mAb, n=9; Tg:mAb+FUS, n=6)

ELISA measurements of Aβ in homogenates from whole mouse hemibrains revealed no significant differences in guanidine-soluble (Tper insoluble) Aβx-42 and Aβx-40 levels between groups (Fig. 3i, j). These results are not surprising given the hippocampal targeting of FUS-mediated BBBD and the short treatment period. However, plaque-lowering in hippocampus suggests that pGlu-Aβ mAb combined with FUS-BBBD exerts additive effects to reduce Aβ plaque deposition in the APP/PS1 the sonicated hippocampus.

### Age-related hippocampal synaptic degeneration is reduced in APP/PS1 mice after treatment with 07/2a and FUS-BBBD

Previous reports have demonstrated that aging in APP/PS1 mice is associated with reductions is synapse number as well as synaptic proteins and mRNA levels in the hippocampus and cortex [28-31]. To determine the effects of FUS and 07/2a on hippocampal synapses in aged APP/PS1 mice, we performed high resolution confocal microscopy analysis [31] of pre- and post-synaptic markers (i.e. synaptic puncta) in hippocampal CA3 in mice treated with mAb alone in combination with FUS-BBBD. The mice in the Tg:mAb+FUS group had significantly more Vglu2-positive (*p* < 0.01), GluR1-positive (*p* < 0.01) and colocalized synaptic puncta (*p* < 0.01) than APP/PS1 mice treated with 07/2a alone (Fig. 4a, b). Western blotting of pre-synaptic markers, Vglut2 and Synapsin-1 (SYN-1), and post-synaptic markers, GluR1 and PSD95, in hippocampal synaptosomes showed significantly elevated synaptic protein levels in mice in the Tg:mAb+FUS group compared to the Tg:PBS or Tg:FUS animals (*p* < 0.01, Fig. 4c, d). The animals in the Tg:mAb group had increased GluR1 compared to the Tg:PBS mice (*p* < 0.01, Fig. 4c, d), but such differences were not found for the other synaptic proteins. WT mice that received PBS only or FUS-BBBD showed significantly higher levels of all 4 synaptic markers compared to the Tg:PBS, Tg:FUS, and Tg:mAb mice (*p* < 0.01), while there was no significant difference in these levels between the Tg:mAb+FUS animals and the WT groups (*p* > 0.05, Fig. 4c, d). These results indicate that APP/PS1 mice with enhanced mAb delivery via FUS-BBBD were at least partially spared synapse loss. This result is in line with our cognitive data and suggests a pro-cognitive health phenotype in the APP/PS1 mice that received the mAb and FUS-BBBD.

**Fig 4.**
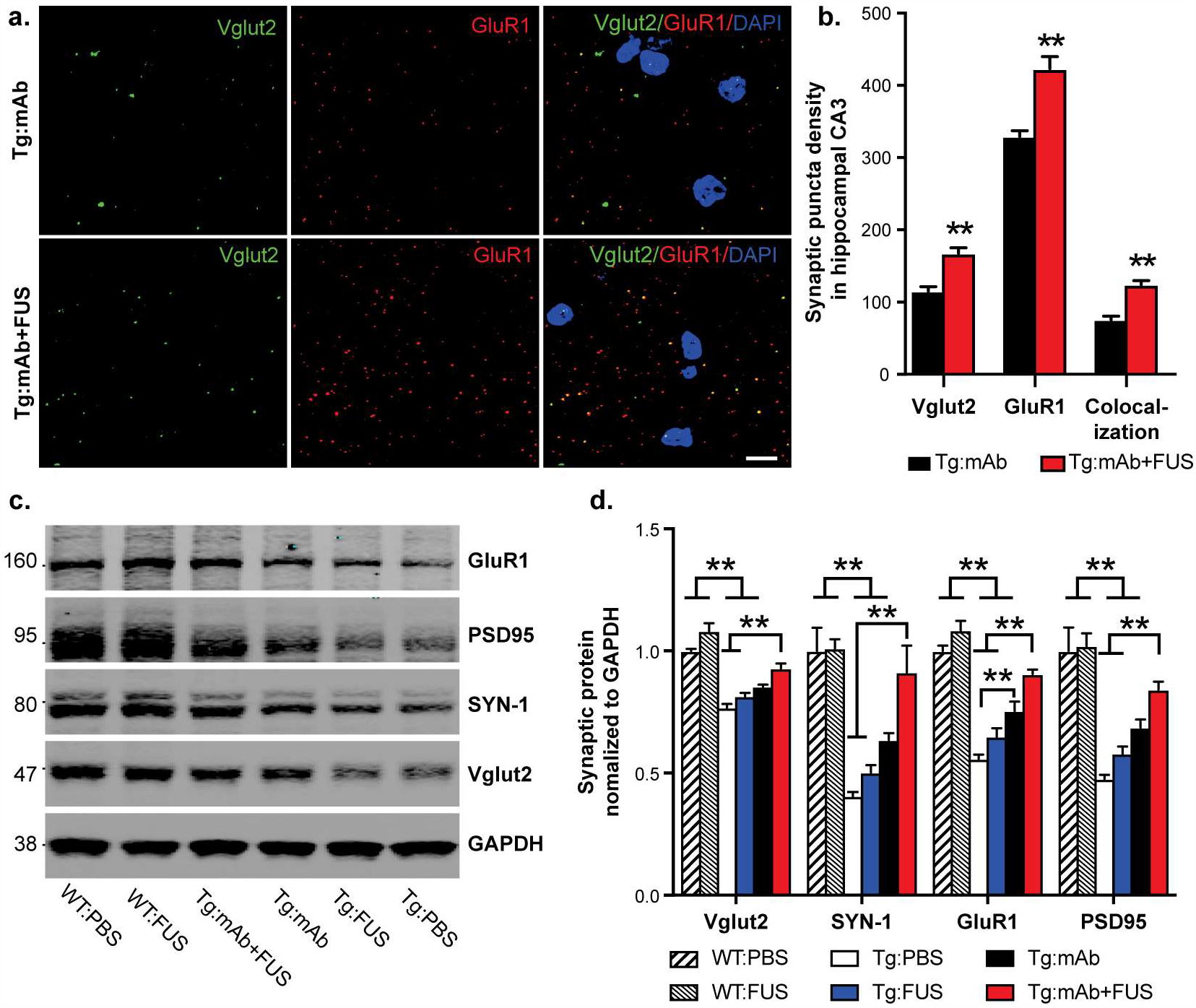
Synaptic puncta analysis and synaptic protein levels in 16 month old mice with or without FUS-BBBD and pGlu3 Aβ mAb. **a**. Synaptic puncta of pre-synaptic and post-synaptic markers Vglut2 and GluR1, respectively, and their colocalizaton in hippocampal CA3 were analyzed by high resolution confocal microscopy in Tg mice treated with mAb alone or in combination with FUS-BBBD. Scale bar = 5 μm. **b**. Mice in the Tg:mAb+FUS group showed increased Vglut1, GluR1 and co-localized synaptic densities compared to the Tg:mAb mice (** *p <* 0.01, 3 equidistant planes, 300 μm apart; one-way ANOVA and Bonferroni post-hoc test). **c**.**d**. Western blotting of synaptic proteins in hippocampal synaptosomes isolated from WT and Tg mice indicated increased levels of the pre-synaptic proteins Vglut2 and SYN-1 and the post-synaptic proteins GluR1 and PSD95 in the Tg:mAb+FUS mice compared to the Tg:PBS and Tg:FUS groups, suggesting a sparing of synaptic loss resulting from the combination of mAb and FUS-BBBD (** *p <* 0.01; WT:PBS, n=13; WT:FUS, n=8; Tg:PBS, n=9; Tg:FUS, n=7; Tg:mAb, n=9; Tg:mAb+FUS, n=6; one-way ANOVA and Bonferroni post-hoc test per marker).

### Altered plaque-associated microglia and macrophage activation in aged APP/PS1 after treatment with 07/2a and FUS-BBBD

Amyloid plaques are often surrounded and infiltrated by immune cells, such as microglia and astrocytes, in AD brain. Thus, we investigated the morphology and colocalization of these cells with plaques in mouse brain by immunostaining for Iba-1 and CD68 (markers for microglia and macrophages) and GFAP (an astrocyte marker). Immunoreactivities of Iba-1 were significantly increased in hippocampal CA3 of the Tg:mAb and Tg:mAb+FUS mice compared to animals in the Tg:PBS group (*p* < 0.05), while no significant difference was found between groups in the pre-frontal cortex (Fig. 5a-c). The Tg:mAb and Tg:mAb+FUS mice had more activated microglia/macrophages associated with Aβ plaque in the hippocampus, as indicated by increased Iba-1 fluorescent intensity (*p* < 0.05, Fig. 5d, e); this result is consistent with reactive microgliosis. In addition, CD68 fluorescent intensity associated with Aβ, indicative of lysosomal phagosomes, was significantly increased over the Tg:PBS animals only in the Tg:mAb group (*p* < 0.05), while no difference was observed in the Tg:mAb+FUS mice (Fig. 5d-f). This result suggests that the pGlu-Aβ mAb, but not FUS-BBBD, induced phagocytosis of Aβ plaques via increased microglia activation or through macrophages. GFAP-positive astrocytes, were clustered around Aβ plaques in the hippocampus of the APP/PS1 mice, but no effects were observed in either the hippocampus or pre-frontal cortex treatment with the anti-pGlu3 Aβ mAb alone or in combination with FUS-BBBD (Fig. 6a, b).

**Fig 5.**
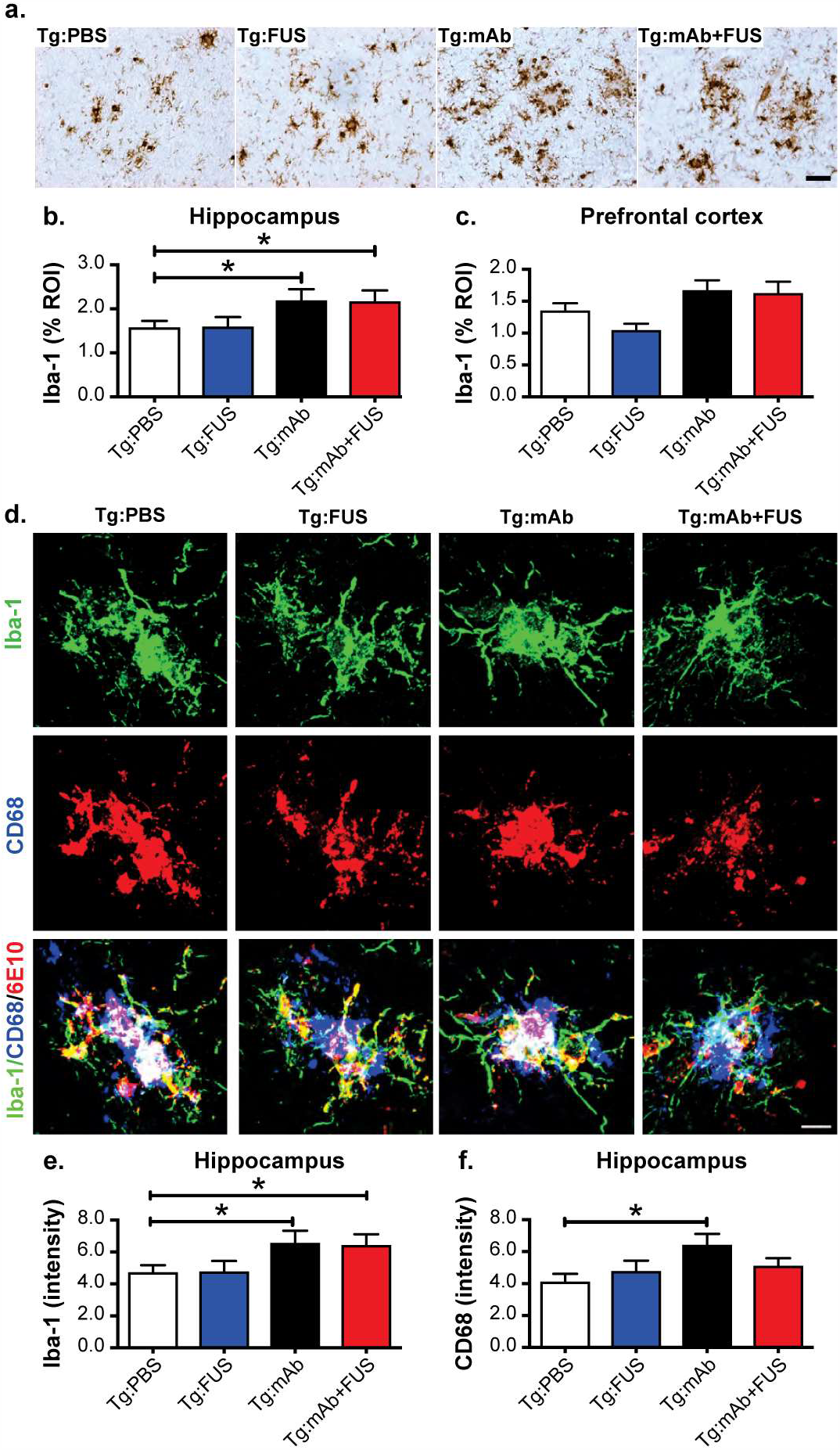
Morphological changes and glial phagocytosis in WT and Tg mice treated with or without FUS-BBBD and pGlu3 Aβ mAb, 07/2a. **a-c**. Iba-1-positive immunostaining (a) showed significant increased activation of microglia and macrophages (i.e. rounder cell bodies and thickened processes) and the clustering of more Iba-1-positive glial cells in sonicated hippocampal CA3 region in the Tg:mAb and Tg:mAb+FUS mice compared to the Tg:PBS group (b) (* *p <* 0.05, 3 equidistant planes 300 μm apart; one-way ANOVA followed by Bonferroni test). Iba-1 positive immunostaining was not significantly different in the non-sonicated prefrontal cortex between groups. Scale bar = 50 μm. **d-f**. High-resolution confocal images of microglia/macrophages (immunoreactive for Iba-1) and phagocytic cells (immunoreactive for CD68) in the hippocampal CA3 region (d), Scale bar = 25 μm. Quantification of immunofluorescence intensity indicated increased microglial acitivation (Iba-1 immunofluoscnent intensity) in the Tg:mAb and Tg:mAb+FUS mice, and increased phagocytosis (CD68 immunofluorescent intensity) in Tg:mAb animals (* *p <* 0.05, compared to Tg:PBS group).

**Fig 6.**
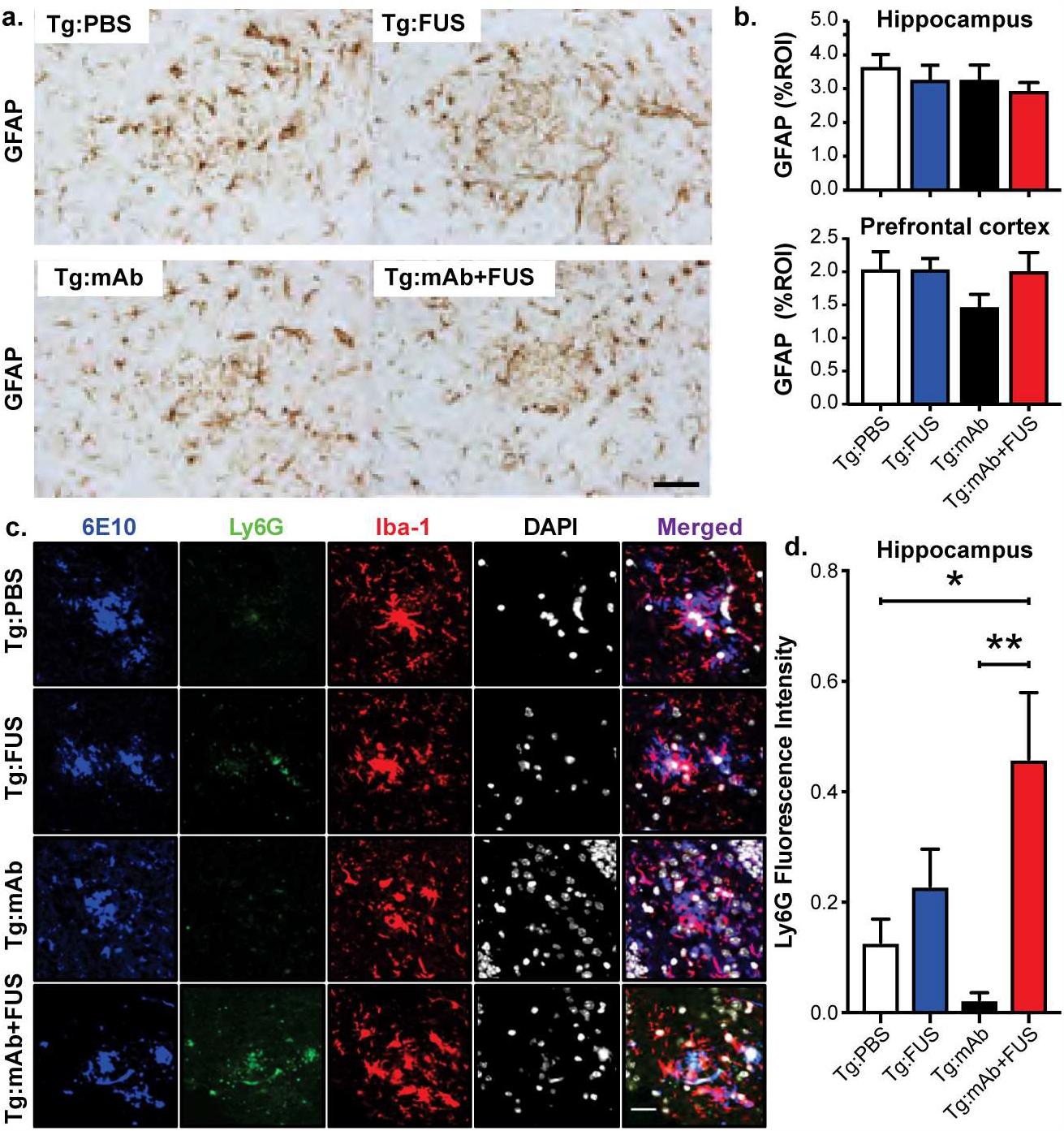
Plaque-associated astroglia and monocytes. **a-b**. Astrocytic immunoreactivity of GFAP in hippocampus and PFC in mice from the Tg:PBS, Tg:FUS, Tg:mAb and Tg:mAb+FUS groups. No significant differences in GFAP immunoreactivity were found between different groups. Scale bar = 50 µm. **c-d**. Confocal microscopy of Ly6G, Iba-1, DAPI, and 6E10 immunofluorescence in hippocampal CA3 of Tg mice with or without FUS-BBBD and pGlu3 Aβ mAb. Scale bar = 25 μm. Fluorescence intensity of Ly6G (b) was significantly higher in the Tg:mAb+FUS group compared to the Tg:PBS and Tg:mAb groups (* *p <* 0.05, ** *p <* 0.01).

### Increased Ly6G-positive plaque-associated immune cells in APP/PS1 mice after treatment with 07/2a and FUS-BBBD

Ablation or inhibition of monocyte migration into the brain has been shown to exacerbate Aβ pathology, while enriching monocytes in blood and enhancing their recruitment to plaque lesion sites greatly diminishes Aβ plaque load in AD transgenic mice [32]. The immunofluorescent intensity of the lymphocyte antigen 6 complex locus G6D (Ly6G), a marker for infiltrating monocytes, showed a significant increase of Ly6G cells associated with Aβ plaques in the Tg:mAb+FUS animals compared to those in the Tg:PBS and Tg:mAb groups (*p* < 0.05, *p* < 0.01, Fig. 6c, d). No significant difference in Ly6G immunofluorescence was found between the Tg:mAb and Tg:PBS groups or between the Tg:FUS group and all other groups. These data suggest that FUS-BBBD in combination with the anti-pGlu3 Aβ mAb facilitated monocyte infiltration into the brain and within the vicinity of the Aβ plaques, which may have contributed to plaque clearance in the mice given the combination treatment.

### FUS-BBBD did not increase microhemorrhage or reduce neuron number

We used hemosiderin staining to investigate the presence of microhemorrhage. With semi-quantification of the number of hemosiderin-positive cells in the whole sagittal brain section, we found FUS-BBBD with or without 07/2a administration did not increase the number of microbleeds in the brain (*p* > 0.05, Fig. 7a,b). These results indicate that the cavitation monitoring and control was successful in ensuring that the FUS treatments did not induce vascular damage or associated damage of brain parenchyma, nor did it enhance microhemorrhages. In addition, we observed no differences between groups in neuronal numbers in hippocampal layers CA1 and CA3, indicating that FUS-BBBD treatment did not affect neurons (Fig. 7c,d).

**Fig 7.**
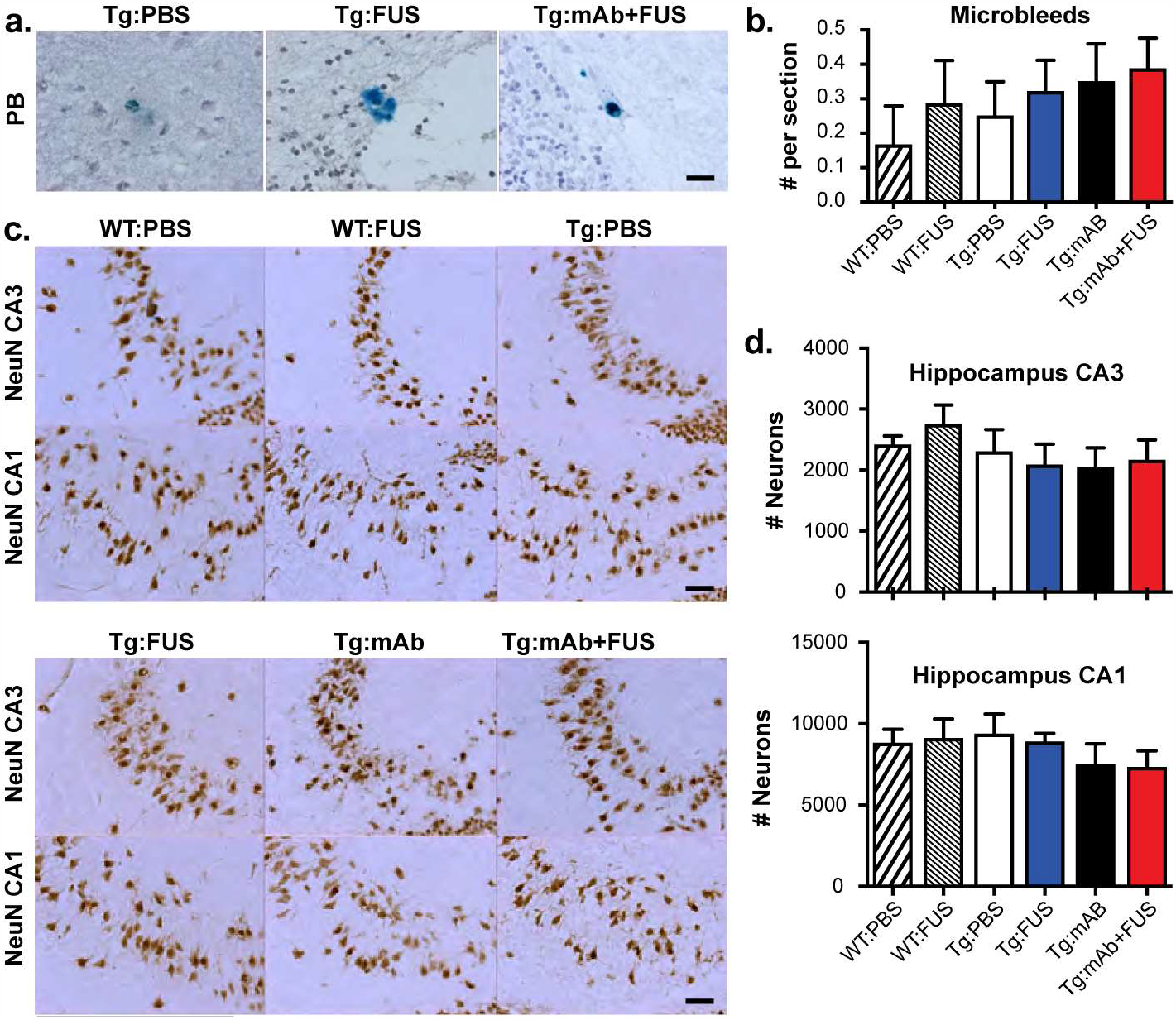
Microhemorrhage and neuron number in WT and Tg mice with or without FUS-BBBD and pGlu3-Aβ mAb. **a-b**. Hemosiderin staining. No significant difference in microhemorrhage number was found between the different groups. Scale bar = 25 μm. **c-d**. Stereology of NeuN-positive cells (indicating neurons) in hippocampal CA3 and CA1. Scale bar = 50 μm. No significant difference in neuron number was found between the different groups.

## Discussion

Our results revealed that anti-pGlu3 Aβ mAb, 07/2a, administered in conjunction with MB-enhanced FUS-BBBD improved learning ability, spared cognitive decline, and preserved synapses in the hippocampus in aged APP/PS1 transgenic mice by engaging microglia and monocytes, thereby promoting clearance of Aβ and reducing the plaque load. A previous study demonstrated that transient BBB opening via FUS and MB induced increased Aβ clearance in AD model mice by microglia activation [15]. Our lab showed that weekly intraperitoneal (i.p.) injections of an anti-pGlu3 Aβ IgG1 mAb (07/1) in young APP/PS1 mice for 28 weeks prevented Aβ plaque deposition and partially spared cognitive performance [21] while i.p. injections of 07/2a in plaque-rich 12 month-old APP/PS1 mice for 16 weeks lowered plaque burden and improved spatial learning and memory in the Water T Maze [22]. Our current results indicate the combination of these two treatments over 3 weeks in aged APP/PS1 mice exerts additive effects compared to either treatment alone, and is in agreement with other work demonstrating that FUS enhances the CNS uptake of peripherally administered antibodies [12, 17] or other therapeutics [13, 33] targeting AD pathology. This work is also in agreement with prior studies in AD model mice [16] and in dogs [34] in that MB-enhanced FUS did not result in increased microhemorrhage or other associated irreversible effects, even in aged animals with significant disease.

Prior work from our lab and others have demonstrated the efficacy of anti-pGlu3 Aβ antibodies in transgenic Alzheimer’s-like mouse models. We reported that starting treatment before and after Aβ deposition resulted significant lowering of pGlu3 Aβ and total Aβ deposits without inducing microhemorrhages in APP/PS1 mice [20-22]. Another study in PDAPP mice demonstrated that a different pGlu3 Aβ specific mAb (mE8-IgG2a; Eli Lilly) also reduced amyloid burden in the absence of microhemorrhages [35]. Our lab has also investigated the effects of anti-pGlu3 Aβ antibodies on cognitive decline in AD-like transgenic mice and reported a significant sparing of spatial learning and memory in the Water T Maze following treatment [19, 22]. Interestingly, plasma Aβ was not elevated in mice treated with the pGlu3 Aβ antibody. In contrast, treatment with 3A1, an N-terminal anti-Aβ mAb, caused a significant rise in plasma Aβ concentration, implying different mechanisms for plaque removal. Currently, a humanized anti-pGlu3 Aβ antibody, Donanemab (LY3002813; Eli Lilly and Co.), is being tested early stage AD patients in clinical trials. The initial results of two small Phase 1 studies, presented at the Alzheimer’s Association International Conference (AAIC, 2016) and Clinical Trials in Alzheimer’s Disease (CTAD, 2019), showed that patients who received a high dose (10-20 mg/kg) of Donanemab by monthly intravenous injections had significant plaque removal in the brain by amyloid PET imaging. Amyloid Related Imaging Abnormalities with Hemorrhage (ARIA-H) were apparent in 2 participants in one trial while the incidence of ARIA-E (Amyloid Related Imaging Abnormalities with Edema), an adverse event associated with amyloid immunotherapy [36], was observed in 1 of every 4 participants in the second Phase 1 trial (www.alzforum.org). While Donanemab was shown to effectively reduce plaques, it induced an anti-drug antibody response in many participants, thereby reducing the antibody’s half-life in the circulation [37, 38]. More recently, Eli Lilly has initiated two larger Phase 2 studies of Donanemab (TRAILBLAZER-ALZ and TRAILBLAZER-ALZ2) in early AD with a primary focus on cognitive and functional outcomes. Eli Lilly presented a press release in mid-January 2021 describing a positive outcome for the TRAILBLAZER-ALZ trial: robust lowering of cerebral amyloid and significant 33% slowing of cognitive and functional decline in patients receiving Donanemab. Our current study suggests that the success of pGlu3 Aβ immunotherapy can be improved with FUS-BBBD without increasing risk for microbleeds.

FUS-BBBD also appears to alter the microenvironment in potentially beneficial ways. Microglia activation and the phagocytosis of Aβ has been observed in anti-Aβ immunotherapy, presumably via microglial Fcγ-receptor binding to antibody-opsonized Aβ aggregates [39]. Our data suggest that anti-pGlu3 Aβ mAb (07/2a) treatment increased the recruitment of microglia to plaques, resulting in increased phagocytosis of Aβ. In addition, infiltrating monocytes were increased in hippocampus after the combined mAb and FUS-BBBD treatment, which appears to have had a beneficial, plaque-clearing effect in this short-term study. Depletion of monocytes or inhibition of monocyte migration into the brain has been shown to worsen Aβ pathology, while enriching monocytes in blood and enhancing their recruitment to plaque lesion sites greatly diminishes Aβ plaque load in AD transgenic mice [32]. It is possible that FUS-BBBD locally modulates activity and migration of microglia and monocytes, accelerating antibody-mediated Aβ clearance in aged AD Tg through increasing cerebral recruitment of monocytes overexpressing Aβ-degrading enzymes, as well as increase peripheral monocyte-derived macrophages.

These changes to the brain we observed after the combined 07/2a + FUS-BBBD treatment may also protect synapses. Consistent with earlier work, we observed plaque-associated hippocampal synaptic degeneration in the aged APP/PS1 mice [40]. However, mice in the Tg:mAb+FUS group had less synapse loss along with a decreased plaque burden. Microglia have been shown to be highly phagocytic and participate in synapse elimination during aging and AD [31, 41]. Perhaps FUS-BBBD facilitates monocyte infiltration and lowers microglia overactivation, thereby reducing microglial synapse phagocytosis and modulating the glial response to pGlu3 Aβ. Further studies are underway to address this mechanistic question.

It has been reported that BBB opening via FUS and MB can markedly ameliorate the pathology of Aβ-deposition and behavior. Burgess et al. found a 19% reduction in plaque number after a single session of FUS-BBBD in 7 month old TgCRND8 mice [14]. Leinenga and Götz found over 50% reductions in both plaque number and area in 12-13 month old APP23 mice after five scanning FUS-BBBD sessions [15]. However, in a subsequent study, that group did not see a reduction in plaque area in 21-22 month old APP23 mice after four scanning sessions, although they did see changes in plaque size and increased numbers of Iba-1-positive cells [16]. Here, in 16 month old APP/PS1 mice, we did not find significant reductions in plaque burden or improvement in learning in the Water T Maze test with three weekly sessions of FUS-BBBD alone, however, some efficacy was seen in contextual fear memory. Overall, these results suggest that treatment with FUS-BBBD alone might be less effective in older mice. However, differences in strain, FUS procedure, and other methodologies might confound direct comparison between studies.

Lastly, our study is limited by the fact that the mice were euthanized 18 days after the last treatment, and therefore, we are unable to quantify antibody delivery to the brain shortly after treatment, when it would be expected to rise. Murine IgG2a antibodies have a half-life of 6-8 days, which may account for the similarity in IgG2a detection in mouse brains between groups at the end of our study. A recent publication from the Aubert lab determined that IVIg delivery into brain was strongly elevated 4 hours post-FUS BBBD treatment, declined moderately by 24 hours and was indistinguishable from sham controls 7 days post-treatment [33]. Therefore, we estimate that the mice that received the combination of 07/2a and FUS BBBD in our study had 3 periods of transient BBB opening that enhanced delivery of 07/2a to the brain. Further studies are underway to better determine 07/2a antibody delivery to brain at different timepoints following FUS-BBBD treatment.

In conclusion, three weekly sessions of MB-enhanced FUS BBBD in the hippocampus combined with i.v. infusion of an anti-pGlu3 Aβ monoclonal antibody selectively reduced plaque burden and synapse loss in the targeted area and improved outcomes in behavior/cognitive tests. The combined procedure did not increase the incidence of microhemorrhage in the context of CAA in aged AD transgenic mice with both plaque and vascular amyloid deposition. Interestingly, our data suggests FUS can ameliorate pathology and neuroinflammation by modulating the activation and infiltration of phagocytes. Therefore, our study highlights the potential of using this technique to improve the brain delivery and thereby, the efficacy of the anti-pGlu3 Aβ mAb.

## Materials and Methods

### Animals

C57BL/6J and APPswe/PS1dE9 breeder mice were obtained from The Jackson Laboratory and bred in-house for this study. Mice were genotyped by PCR using the following primers: APP/PS1: 5’-GACTGACCACTC GACCAGCTT-3’ and 5’-CTTGTAAGTTGGATTCTCATAT-3’; Mice were aged to 16 months. Only males were used for this study to reduce gender-specific variability in the behavioral tests. At the end of the study, mice were anesthetized, blood collected, and the brain perfused with saline prior to harvest.

All animal studies were approved by the Institutional Animal Care and Use Committees at Harvard Medical School and Brigham and Women’s Hospital. The Harvard Medical School and Brigham and Women’s Hospital animal care and management programs are accredited by the Association for the Assessment and Accreditation of Laboratory Animal Care, International (AAALAC) and meet the National Institutes of Health standards as set forth in the 8^th^ edition of the Guide for the Care and Use of Laboratory Animals. The institutions also accept as mandatory the PHS Policy on Humane Care and Use of Laboratory Animals by Awardee Institutions and NIH Principals for the Utilization and Care of Vertebrate Animals Used in Testing Research and Training. There is on file with the Office of Laboratory Animal Welfare (OLAW) an approved Assurance of Compliance: A3431-01 for Harvard Medical School and A4752-01 for Brigham and Women’s Hospital.

Animals were anesthetized by i.p. injections of ketamine (80 mL/kg/h) and xylazine (10 mL/kg/h) or isoflurane before FUS-BBBD or administration of mAb or PBS. Isoflurane was applied through a nosecone with medical air as the carrier gas. The isoflurane concentration was titrated based on the respiration rate and was typically 1-2%. A catheter was placed in the tail vein for i.v. administration.

### Antibody, FUS-BBBD, and/or PBS Treatment

A total of 52 male mice were utilized in this study (∼23-35 g). Sixteen month-old mice were divided into 6 groups and received one of the following treatments: WT:PBS, 130 μl sterile PBS i.v. in WT mice (n = 13); WT:FUS, FUS-BBBD in WT mice (n = 8); Tg:PBS, 130 μl sterile PBS i.v. in APP/PS1dE9 mice (n = 9); Tg:FUS, FUS-BBBD in APP/PS1dE9 mice (n = 7); Tg:mAb, 500 μg anti-pGlu3 Aβ 07/2a mouse monoclonal antibody (divided into 4 consecutive infusions of 13 μl each containing 125 µg mAb) generated and provided by Vivoryon Therapeutics AG (Halle, Germany), i.v. in APP/PS1dE9 mice (n = 9); Tg:mAb+FUS, 500 μg anti-pGlu3 Aβ 07/2a mouse monoclonal antibody (13 μl x 4) immediately prior to FUS-BBBD in APP/PS1dE9 mice (n = 6). Mice were treated with a total volume of 130 μl liquid (antibody and/or PBS and/or MB) to limit the volume of liquid that mouse can tolerate. Behavior tests were initiated one week after the third and final treatment.

### FUS-BBBD

Before the ultrasound exposures, the scalp was shaved and remaining hair on the head was removed using depilatory cream to ensure unimpeded ultrasound propagation. A catheter was placed in the tail vein, and the animal’s head was fixed in a plastic stereotactic frame that was built in-house. This frame was placed on the FUS device. Sonications (10-ms bursts applied at 2 Hz for 100 s) were applied coincident with an injection of the MB ultrasound contrast agent Optison (GE Healthcare, Little Chalfont, Buckinghamshire, UK; dose: 100 µl/kg; diluted 4x in PBS) to open the BBB. Two locations in each hemisphere were sonicated in each session (± 2 mm lateral, 3 mm dorsal, and 2 and 3.5 mm anterior to the interaural line). The time between sonications was at least two minutes to allow for the MB to mostly clear from circulation.

A FUS system with cavitation-controlled transmission was designed and built in-house [42]. An air-backed, spherically curved transducer (diameter/radius of curvature: 10/8 cm) was used for FUS transmission with a resonant frequency of 278 kHz. The transducer was attached to a three-axis positioning system (A4000, Velmex, Bloomfield, NY, USA) (Fig. 1a). The transcranial sonications were applied at the third harmonic of the FUS transducer (835 kHz) via a function generator (33220A, Agilent, Santa Clara, CA, USA) and an amplifier (240L, E&I, Rochester, NY, USA). The electrical power was measured using a power meter (E4419B, Agilent, Santa Clara, CA, USA) and dual-directional coupler (C5948-10, Werlatone, Patterson, NY, USA); a radiation force balance was used to measure the transducer efficiency. The pressure field in the focal plane was mapped using a needle hydrophone (HNC-1000; Onda, Sunnyvale, CA). This map along with radiation force balance measurements were used to estimate the pressure amplitude at the focus in water. The half-width and -length of the 50% isopressure contours were 1.9 and 11.4 mm respectively. The pressure amplitude used in the mouse sonications was 0.33 MPa (estimate in water).

Acoustic emissions were monitored using a 10-element piezoelectric hydrophone. The elements of this passive cavitation detector were arranged in a ring (inner/outer diameter: 13/15 cm) that surrounded the transducer. The cavitation emission signals were recorded with a 12-bit high-speed digitizing card (PCX-5124, National Instruments, Austin, TX, USA). Frequency spectra were generated via fast Fourier transform, and the harmonic and wideband emissions were tracked in real time using software developed in-house [42] in MATLAB (MathWorks, Natick, MA, USA). The software acquired baseline emissions prior microbubble injections and then waited for the user to administer microbubbles. Once the injection started, harmonic (2×, 3×, 4× 835 kHz) and wideband emissions were calculated in dB relative to baseline emissions. This cavitation-controlled system was used to confirm that stable cavitation was occurring (detected by harmonic emission), as well as to avoid inertial cavitation (assessed by wideband emission). If wideband emission was detected in the cavitation controller, the exposure level was automatically decreased.

To validate this system, a pilot study was performed in WT mice that visualized the BBB opening by obtaining fluorescence imaging of the delivery of Trypan blue after sonication. The images were obtained with a system developed in-house consisting of two red LED lamps (610-630 nm, GR-PAR38-12W-R-1, ABI, Indianapolis, IN) for excitation, and a bandpass filter (687-748 nm, 715AF58, Omega Optical, Brattleboro, VT, USA) and digital camera (C920, Logitech) for image acquisition.

### Behavior tests (performed in the following order)

#### Open Field Test (OF)

The OF test was used to assess general locomotor activity in the mouse. The total distance traveled (cm) was used as a measure of general locomotor activity, while the percent distance traveled in the center was used to measure anxiolytic behavior. Methods were performed as previously described [43].

#### Water T Maze Test (WTM)

The WTM test examines spatial learning and memory by training mice to use spatial cues in a room to navigate to a hidden platform to escape water. We performed the test as previously reported [43].

#### Contextual Fear Conditioning Test (CFC)

Contextual Fear Conditioning was used to assess fear learning and memory as mice tend to show a fear response (freezing) when exposed to an aversive brief foot shock stimulus. Learning is assessed during training on Day 1 whereas memory is assessed by the time spent freezing when the mice are re-exposed to the same context on Day 2 in the absence of a foot shock. Testing was performed as previously reported [43].

### Pathological and Biochemical Analyses

#### Immunohistochemistry

Mouse hemibrains were fixed in 4% PFA for 24 h, cut into 10 µm cryosections, mounted onto microscope slides, and immunostained as previously described [43]. Brain sections were incubated with anti-Aβx-42 (1:200, Covance), anti-Aβ 6E10 (1:1000, Covance), anti-pGlu3 IgG2b (1µg/mL; Vivoryon Therapeutics) and NeuN (1:200, Serotec) mouse monoclonal antibodies, or Aβ40 (1:200, Covance), Iba-1 (1:200, Wako) and GFAP (1:1000, DAKO) rabbit polyclonal antibodies, DAPI or CD68 (1:250, Serotec) rat polyclonal antibody overnight at 4°C. Sections were washed in TBS and then incubated with biotinylated secondary antibodies and developed using Vector ELITE ABC kits (Vector Laboratories) and 3,3-diaminobenzidine (Sigma-Aldrich) or immunofluorescent-labeled secondary antibodies and cover-slipped with mounting media (Vector).

#### Aβ load analysis

Aβ immunoreactivity for all mice was captured by imaging sections in a single session under a Nikon Eclipse E400 microscope. Aβ plaque load and Aβ plaque size were quantified within a region of interest (ROI) using the Bioquant image analysis system (Nashville, TN). Three sections at equidistant planes were analyzed per mouse by an operator who was blinded to mouse genotype and treatment.

#### Aβ and cytokine ELISAs

Tper-insoluble, guanidine hydrochloride-extracted brain homogenates were extracted from mouse hemibrain, including cortex and hippocampus, and run on an MSD Aβ Triplex ELISA as previously described [43].

#### Confocal analysis of glia morphology and association with Aβ plaques

Immunofluorescence for Aβ plaques (6E10), microglia/macrophages (Iba-1) and (CD68) was detected by confocal microscopy (Zeiss, LSM 710; Carl Zeiss). Images were collected using the same exposure settings. Confocal Z-stack images (optical slices of 0.2 µm) of plaques and surrounding glia were collected using a 63x objective. Three images were acquired from 3 equidistant planes, 500 μm apart per mouse. Immunofluorescent intensity analysis and 3D reconstruction of Z-stack images were performed with confocal image analysis software, Zein black (Carl Zeiss). Glia counts were performed using stereological methods after collecting images.

#### Synaptic puncta staining and analysis was performed as previous described [43, 44]

Confocal imaging was performed using a ZEISS LSM710 confocal microscope and a 63x oil objective. Images were acquired using a 1 Airy Unit (AU) pinhole, while holding constant the gain and offset parameters for all sections and mice per experiment.

#### Preparation of synaptosome fractions

Hippocampal synaptosome fractions were prepared as described previously [43, 45].

#### Western blotting of synapse markers

Western blotting performed on hippocampal synaptosomes as previously described [31, 43] using rabbit polyclonal antibody Synapsin-1 (SYN-1, 1:200; Millipore), goat anti-guinea pig antibody VGlu2 (1:1000; Millipore), goat anti-rabbit antibody GluR1 (1:200; Abcam) and mouse monoclonal antibody PSD95 (1:200; Millipore), and GAPDH (1:200; Millipore). Blots were scanned using a LiCor Odyssey Infrared Imaging System. Intensity of bands was measured by LiCor Odyssey software.

#### Stereological quantification of neurons

Immunohistochemistry for anti-neuronal nuclear protein, NeuN (1:250; Millipore Bioscience Research Reagents) was performed as described previously [46]. Stereological counting of neurons and glia was performed on 10 μm sagittal, immunostained stained sections at each of 6 planes 250μ m apart per mouse using the optical dissector method. Cells were counted in an area of ∼300 µm within a 10 µm depth per brain region examined. The number of neurons and glia per section in each brain region was estimated using ∼15 optical dissectors and the Bioquant image analysis system according to the principles of Cavalieri [47].

### Statistics

#### Statistical analysis

Data are expressed as mean ± SEM. Comparisons were made between WT:PBS vs. WT:FUS, WT:PBS vs. Tg:PBS, Tg:PBS vs. Tg:mAb, Tg:PBS vs. Tg:FUS, Tg:PBS vs. Tg:mAb+FUS, Tg:mAb vs. Tg:mAb+FUS, and Tg:FUS vs. Tg:mAb+FUS groups. Significance for all behavioral tests was assessed by one-way ANOVA followed by Fisher’s Protected Least Significant Difference using StatView Version 5.0 software. All other data were analyzed by one-way or two-way ANOVAs followed by Bonferroni’s post hoc test or the Student *t*-test using Prism Version 6.0 (GraphPad) software. A *p* value of *<* 0.05 was considered significant.

## Acknowledgements

The authors thank Grace Liu (BWH Neurology) for her assistance with mouse breeding and P. Jason White and Can Barış Top (BWH Radiology) for their assistance with the experiments. We thank Jens-Ulrich Rahfeld for critical comments on the manuscript. Vivoryon Therapeutics is thanked for providing the 07/2a mAb as a gift-in-kind.

## Funding

This work was funded by grants from The Focused Ultrasound Foundation (Charlotteville, VA) to CAL and NJM, the National Institutes of Health, NIH/NIA R01 AG040092 and NIH/NIA RF1 AG058657 to CAL, and NIH/NIBIB R01 EB028686 to NJM.

## Availability of Data and Materials

The datasets generated and/or analyzed during the current study are available from the corresponding authors upon reasonable request.

## Authors’ Contributions

Qiaoqiao Shi: Investigation, Methodology, Roles/Writing - original draft, Writing - review & editing. Tao Sun: Investigation, Methodology, Writing - review & editing. Yongzhi Zhang: Investigation, Writing - review & editing. Chanikarn Power: Investigation, Writing - review & editing. Camilla Hoesch: Investigation, Methodology. Shawna Antonelli: Investigation, Methodology. Maren K. Schroeder: Investigation, Methodology, Writing – review and editing. Barbara J. Caldarone: Formal analysis, Investigation, Methodology, Writing - review & editing. Nadine Taudte: Investigation, Methodology. Mathias Schenk: Investigation, Methodology. Thore Hettmann: Writing - review & editing. Stephan Schilling: Conceptualization, Investigation, Resources, Writing - review & editing. Nathan J. McDannold: Conceptualization, Data curation, Funding acquisition, Investigation, Methodology, Project administration, Resources, Software, Supervision, Writing - review & editing. Cynthia A. Lemere: Conceptualization, Data curation, Funding acquisition, Investigation, Methodology, Project administration, Resources, Supervision, Roles/Writing - original draft, Writing - review & editing.

## Declaration of Interest

We declare that Stephan Schilling is a former and Thore Hettmann, present, employees of Vivoryon Therapeutics N.V., Germany, and hold stock options of the company. Stephan Schilling is an advisor to Vivoryon Therapeutics N.V. Co-senior author, Cynthia A. Lemere, was an unpaid scientific advisory board member for Vivoryon Therapeutics N.V., receives antibodies, and has previously received unrestricted funding from Vivoryon Therapeutics N.V. for some of her past work on pGlu3 Aβ immunotherapy. She serves as a consultant to Biogen and Acumen Pharmaceticals, both of which are developing anti-amyloid immunotherapies. Brigham and Women’s Hospital holds two patents related to this work.

